# BLMPred: predicting linear B-cell epitopes using pre-trained protein language models and machine learning

**DOI:** 10.1101/2025.09.10.675202

**Authors:** Barnali Das, Dmitrij Frishman

## Abstract

B-cells get activated through interaction with B-cell epitopes, a specific portion of the antigen. Identification of B-cell epitopes is crucial for a wide range of clinical applications, including disease diagnostics, vaccine and antibody development, and immunotherapy. While experimental B-cell epitope identification is expensive and time-consuming, computational tools are starting to emerge that can generate lists of high-confidence epitopes for experimental trials. In this paper, we present BLMPred, a sequence-based linear B-cell epitope prediction tool, which exploits pretrained protein Language Model embeddings for deriving local and global protein structural features from the protein primary structure. BLMPred is a binary classifier which can predict whether an input peptide sequence is an antibody epitope or not without relying on 3D protein structures. BLMPred has been shown to outperform other comparable tools when tested on multiple independent datasets. It is freely available at https://github.com/bdbarnalidas/BLMPred.git.

## 1 Introduction

B-cells provide long-term immunity against cancerous cells and pathogens/antigens and hence, are a vital component of the adaptive immune system. B-cells get activated when B-cell receptors, transmembrane proteins present on the B-cell surface, interact with B-cell epitopes, a specific portion of the antigen. Linear B-cell epitopes are stretches of adjacent residues along the antigen primary sequence, whereas non-contiguous residues spatially co-localized by protein folding form discontinuous or conformational B-cell epitopes. Both linear and discontinuous B-cell epitopes play significant roles in binding with immunoglobulins (B-cell receptors or antibodies) for recognizing foreign antigens (da Silva, Ascher et al. 2023).

Identification of B-cell epitopes is crucial in clinical and biotechnological applications such as disease diagnostics (Mucci, Carmona et al. 2017), vaccine and antibody design (Kozlova, Cerf et al. 2018, Behmard, Soleymani et al. 2020). Since B-cell epitope identification by assay screening is expensive and time-consuming, computational tools are nowadays becoming a major requirement for reducing development time and cost (Shirai, Prades et al. 2014, da Silva, Ascher et al. 2023).

The accuracy of in-silico B-cell epitope prediction methods has seen significant improvements over the past decades. Available approaches can be broadly classified into two categories. The first category comprises methods accepting protein sequences as input, such as Bepipred-3.0 (Clifford, Høie et al. 2022), Bepipred-2.0 (Jespersen, Peters et al. 2017), epiDope (Collatz, Mock et al. 2020), GraphBepi (Zeng, Wei et al. 2023), Emini surface accessibility prediction (Emini, Hughes et al. 1985), Parker hydrophilicity prediction (Parker, Guo et al. 1986), Kolaskar & Tongaonkar antigenicity prediction (Kolaskar and Tongaonkar 1990), Bepitope (Odorico and Pellequer 2003), and BcePred (Saha and Raghava 2004). The second category includes tools that have been trained explicitly on exact linear B-cell epitope sequences, such as SVMTriP (Yao, Zheng et al. 2020), LBEEP (Saravanan and Gautham 2015), epitope1D (da Silva, Ascher et al. 2023). These tools are beneficial when the user wants to know whether the input peptide may be a potential linear B-cell epitope or not, rather than recognizing the potential B-cell epitope within the given protein sequence. Methods from the first group seem to perform poorly when tested on short peptides, as observed in a recent study (da Silva, Ascher et al. 2023).

Here, we propose BLMPred for predicting whether an input peptide sequence in the length range of 5-60 is a linear B-cell epitope or not. BLMPred was developed by training a Support Vector Machine on the numerical embeddings generated by the ProtTrans protein Language Model (pLM) from a huge experimentally curated linear epitope dataset. On a comprehensive benchmark dataset BLMPred outperformed SVMTriP, LBEEP, and epitope1D based on most performance metrics.

## 2 Materials and methods

### 2.1 Dataset of linear B-cell epitopes

We downloaded 208265 experimentally validated linear B-cell epitope sequences (positive samples) and 487127 non-B-cell epitope sequences (negative samples) from the Immune Epitope Database (IEDB, version: March 2023) (Vita, Mahajan et al. 2019). We removed 1075 duplicate entries, 215 peptides containing non-standard amino acids symbols (Z, B, J, O, U, X), 512 peptides with a length less than the typical minimum length of 5 amino acids for linear B-cell epitopes (De and Tomar 2014), as well as 95971 peptides found both in the positive and the negative dataset. Furthermore, all peptides in the positive dataset longer than 60 amino acids were excluded from consideration since the negative dataset contained peptides with the lengths of up to 60 amino acids. This dataset was named BLMPred_5to60.

Recent reports suggest that the length of B-cell epitopes varies between 5-8 and 25 amino acid residues (Manavalan, Govindaraj et al. 2018, Ashford, Reis-Cunha et al. 2021, Ras-Carmona, Lehmann et al. 2022, da Silva, Ascher et al. 2023). This length range is dictated by the structural requirements of epitope binding to the Complementarity Determining Regions (CDRs) of the B-cell receptors. According to the INDI database (Deszyński, Młokosiewicz et al. 2022), CDR1, CDR2, and CDR3 vary in lengths between 4-17, 5-17, and 5-38 amino acids, respectively. We therefore created an alternative dataset, BLMPred_8to25, consisting of peptides varying in length between 8 and 25. After data cleaning steps, BLMPred_5to60 and BLMPred_8to25 contained 111015 (390589) and 102023 (387155) positive (negative) samples, respectively.

### 2.2 Preparation of training and test datasets

Since the BLMPred_5to60 and BLMPred_8to25 datasets were initially imbalanced, with the positive to negative sample ratio of 1:3, we made them balanced by drawing 111015 and 102023 negative samples from the pool of 390589 and 387155 negative samples in these two datasets, respectively. Additionally, we ensured that the sequence length distribution of these negative samples closely matched that of the positive samples. Both datasets were split into training (90%) and test (10%) datasets while retaining similar length distributions (Supplementary Figure S1). The training and test datasets constructed from BLMPred_5to60 (BLMPred_8to25) datasets are referred to as BLMPred_5to60_training (BLMPred_8to25_training) and BLMPred_5to60_test (BLMPred_8to25_test), respectively. The training datasets were used for cross-validation while the test datasets were solely used as independent datasets for assessing the performance of the final trained methods.

### 2.3 Preparation of benchmarking dataset

We prepared a separate independent dataset (BLMPred_benchmark) for comparing the performance of BLMPred with other existing tools. Since the training and test datasets described above are based on the IEDB release of March 2023, we downloaded sequences of linear B-cell epitopes and non-B-cell epitopes deposited with IEDB after April 2023. After data cleaning and filtering, the final BLMPred_benchmark dataset contained 2928 positive and 1000 negative samples.

### 2.4 Language Model (LM) embeddings

For each peptide in our dataset, we generated average embeddings of length 1024 by utilizing the ProtT5-XL-U50 model of the ProtTrans Protein Language Model (Elnaggar, Heinzinger et al. 2021).

### 2.5 Machine learning

Identification of linear B-cell epitopes was cast as a binary classification problem where an input peptide was either a B-cell epitope or not. A broad range of traditional machine learning models implemented in the Scikit-learn package (Pedregosa, Varoquaux et al. 2011) was tested, including adaboost classifier, bagging classifier, extra trees classifier, Gaussian Naïve Bayes, histogram-based gradient boosting classifier, k-nearest neighbors, linear discriminant analysis, logistic regression, multi-layer perceptron, quadratic discriminant analysis, random forest, and support vector machine. Also, we tested the XGBoost classifier (Chen and Guestrin 2016) from the XGBoost package on the Python language platform. Furthermore, we trained an Explainable Boosting Machine (EBM) learning (Nori, Caruana et al. 2021) classifier provided by an open source Python package, InterpretML (Nori, Jenkins et al. 2019). We utilized RAPIDS, a datascience framework capable of executing end-to-end pipelines completely in the GPU (Hricik, Bader et al. 2020) for an accelerated training of the machine learning models.

### 2.6 Performance evaluation metrics

To assess the model performance on the test dataset, we calculated several performance metrics including accuracy (ACC), precision (P), sensitivity or recall (R), F1 score (F1), specificity (S), Matthew’s Correlation Coefficient (MCC), area under the ROC curve (AUROC), and the area under the precision-recall curve (AUPRC) as follows:

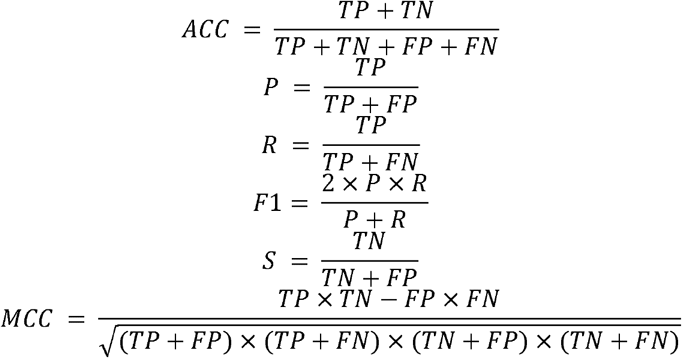

where TP, TN, FP, and FN denote the number of true positives, true negatives, false positives, and false negatives, respectively. Among all the evaluation metrics, MCC has been reported to be more informative in evaluating binary classification problems (Chicco 2017, Chicco and Jurman 2020). Hence, although we report our results based on the full set of metrics, we selected the trained model with the highest MCC measure as the optimal classifier for further processing.

## 3 Results and discussions

### 3.1 Overview of the methodology

A high-quality dataset of experimentally verified linear B-cell epitopes (BLMPred) was derived while keeping similar length distributions of peptides among the positive and negative samples. Peptide sequences were converted into feature vectors of 1024 numerical values using the ProtTrans embedder. The generalization capability of models was first evaluated on the training data based on 10-fold cross-validation, while the final accuracy assessment was conducted on test data. We trained a broad range of machine learning models and selected the best performing model, which was then applied to predict linear B-cell epitopes in the independent dataset. The entire methodology has been summarized in Figure 1.

**Figure 1.**
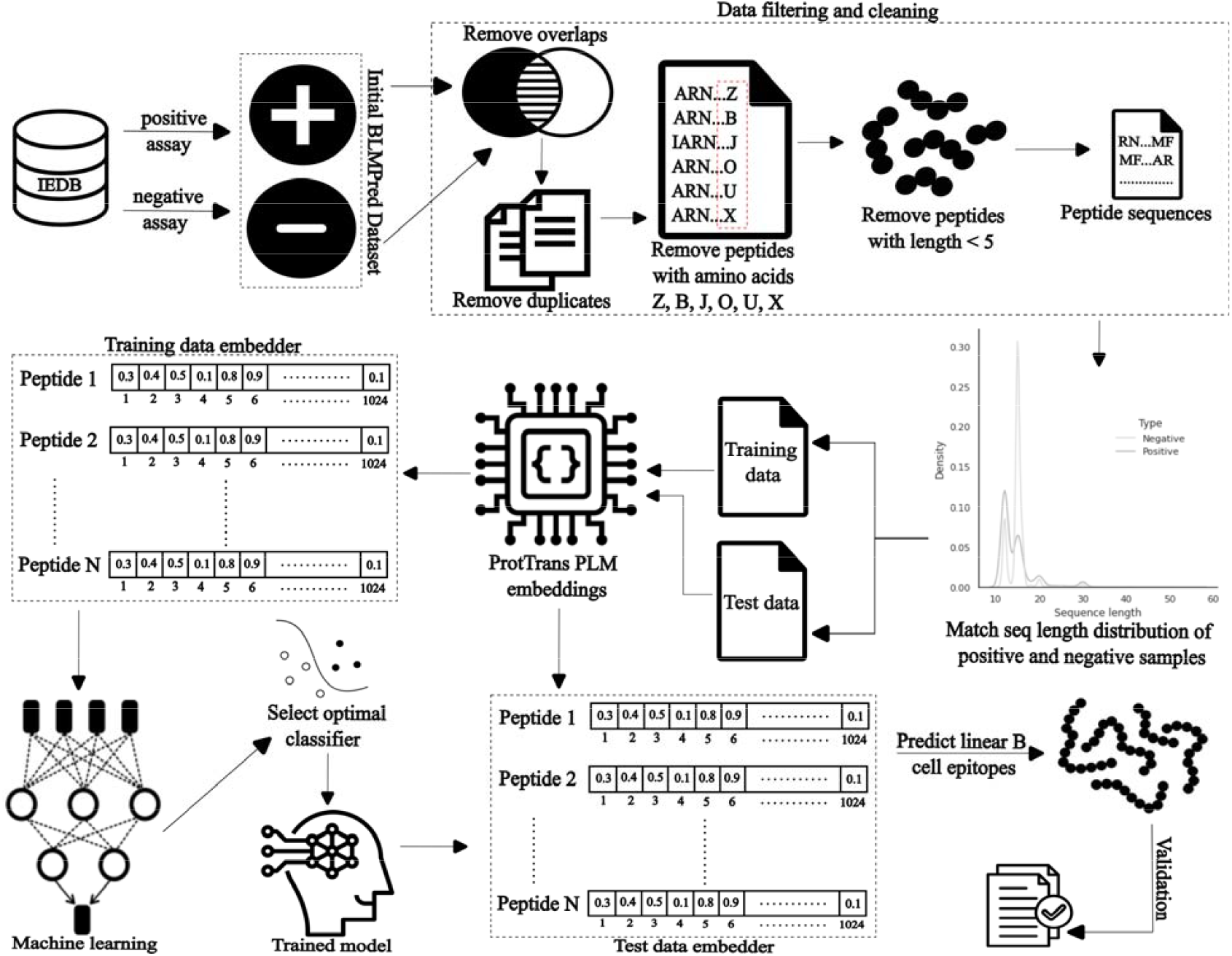
Flowchart of our proposed methodology

### 3.2 Models trained on the BLMPred_5to60 dataset

Machine learning models listed in Section 2.4 were trained on the embeddings derived from the BLMPred_5to60 dataset and their performance was assessed by 10-fold cross validation (Supplementary Table S1, Supplementary Figure S2). The best results were achieved with Support Vector Machine, with the mean±std values of accuracy, precision, recall, F1 score, specificity, MCC, AUROC and AUPRC of 0.839±0.0027, 0.849±0.0046, 0.824±0.0037, 0.837±0.0033, 0.855±0.0041, 0.679±0.0055, 0.839±0.0028, and 0.788±0.0047, respectively. For each classifier, the 10 trained models resulting from each fold of the 10-fold cross-validation were tested on the independent BLMPred_5to60_test dataset (Supplementary Table S2, Supplementary Figure S3) and SVM outperformed all other classifiers. Thus, when utilizing ProtTrans embeddings as input, SVM was clearly the best performing method on the BLMPred_5to60 dataset. In general, we found that none of the models were overfitted and their results were quite robust, as evidenced by the low standard deviation values of the performance metrics (Supplementary Tables S1 and S2).

### 3.3 Models trained on the BLMPred_8to25 dataset

SVM was also the best performing method among the machine learning models trained on the embeddings derived from the BLMPred_8to25 dataset (Supplementary Tables S3, S4, Supplementary Figures S4, S5). Its accuracy was around 5% higher than that of the next best performing model, k-nearest neighbors. SVMs trained on the BLMPred_8to25 and BLMPred_5to60 datasets achieved a similar performance (Supplementary Tables S1-S4). This is not surprising as the BLMPred_8to25 and BLMPred_5to60 strongly overlap: only approximately 9% of the BLMPred_5to60 dataset is constituted by B-cell epitopes with lengths outside of the 8-25 range which have been eliminated to create the BLMPred_8to25 dataset.

### 3.4 BLMPred models

Based on the model assessment presented above, we selected SVM for further analyses. SVM trained on the entire BLMPred_5to60_training and BLMPred_8to25_training datasets will be referred to as BLMPred_5to60 and BLMPred_8to25 models, respectively. The BLMPred_5to60 model, when tested on the independent BLMPred_5to60_test dataset, exhibited the accuracy, precision, recall, F1 score, specificity, MCC, AUROC and AUPRC of 0.846, 0.859, 0.829, 0.844, 0.864, 0.693, 0.846, and 0.798, respectively. Similarly, the values of accuracy, precision, recall, F1 score, specificity, MCC, AUROC and AUPRC achieved by the BLMPred_8to25 model when tested on the independent BLMPred_8to25_test dataset were 0.835, 0.849, 0.814, 0.831, 0.856, 0.671, 0.835, and 0.785, respectively. The BLMPred_5to60 and BLMPred_8to25 models accurately predicted 82.9% (86.4%) and 81.4% (85.6%) of the B-cell epitopes (non-B-cell epitopes) present in the BLMPred_5to60_test and BLMPred_8to25_test datasets, respectively, with low Type I and Type II error levels (Figure 2).

**Figure 2.**
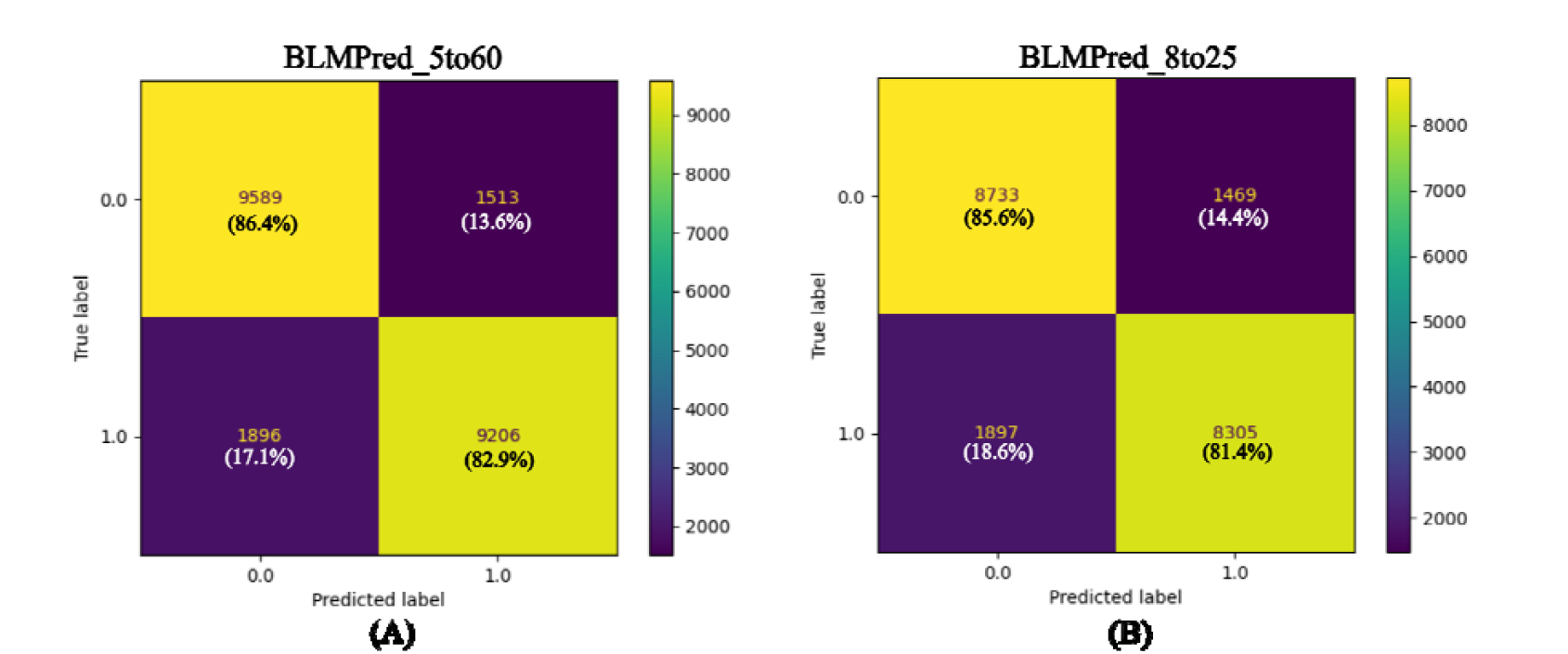
Confusion matrices obtained (A) on the independent BLMPred_5to60_test dataset by the BLMPred_5to60 model, and (B) on the BLMPred_8to25_test dataset by the BLMPred_8to25 model.

### 3.5 BLMPred performance on homology reduced dataset

Strict homology reduction procedures routinely applied to full-length protein sequences are not directly applicable to short peptides. Nevertheless, we assessed potential similarity between the peptides in our datasets as well as model performance on a homology reduced dataset. We downloaded 215912 experimentally validated linear B-cell epitope sequences (positive samples) and 491874 non-B-cell epitope sequences (negative samples) from the Immune Epitope Database (IEDB, version: November 2024) (Vita, Mahajan et al. 2019). The B-cell epitopes and non-B-cell epitopes belonged to 10155 and 7925 host proteins, respectively, with unique UniProt accession numbers. Protein sequences of these host proteins in FASTA format were extracted using Dbfetch (Madeira, Pearce et al. 2022) and clustered based on 70% sequence similarity by CD-HIT (Huang, Niu et al. 2010). To reduce redundancy, training and test datasets were generated by extracting epitopes from the representative sequences of each cluster. After extensive data cleaning and filtering, the final dataset comprised epitope sequences with lengths varying between 5-60. Since the datasets are imbalanced (negative to positive ratio 4:1), we balanced them by drawing samples from the negative dataset while keeping a similar sequence length distribution as that of the positive samples. The final training and testing datasets consisted of 115310 and 20926 epitopes, respectively. ProtTrans embeddings (Elnaggar, Heinzinger et al. 2021) of these datasets were generated and used for training machine learning models. Support Vector Machine performed best among all models when tested on the test dataset and achieved the accuracy, precision, recall, F1 score, specificity, MCC, AUROC and AUPRC of 0.754, 0.92, 0.555, 0.693, 0.952, 0.553, 0.754, and 0.734, respectively.

### 3.6 Performance of BLMPred compared with the reported performance of other linear B-cell epitope prediction tools

We compared the performance of BLMPred with three other models trained to classify an input peptide sequence as being a B-cell epitope or not: Support Vector Machine based on Tri-peptide similarity and Propensity scores (SVMTriP) (Yao, Zheng et al. 2020), Linear B-Cell Exact Epitope Predictor (LBEEP) (Saravanan and Gautham 2015), and epitope1D (da Silva, Ascher et al. 2023). A detailed summary of these tools is presented in Supplementary Table S5, including the specific algorithms, datasets and features utilized as well as the performance metrics reported in the corresponding original publications. According to Supplementary Table S5, epitope1D reportedly outperforms other methods in terms of MCC and AUROC, while our method, BLMPred_5to60, performs better than all existing tools in terms of accuracy, precision, recall, specificity, F1 score, and AUPRC. We attribute this high-performance level of BLMPred_5to60 to many up-to-date experimentally verified linear B-cell epitope data collected from IEDB, extensive data filtering, and the utilization of ProtTrans embeddings. In the next section, we utilize a benchmarking dataset and perform a detailed comparative analysis of BLMPred with the SVMTriP, LBEEP, and epitope1D.

### 3.7 BLMPred compared with other existing tools on an independent dataset

As mentioned in Section 3.5, there are two major varieties of B-cell epitope prediction tools: i) methods predicting epitopic regions from protein sequences, and ii) methods predicting the possibility of an input peptide to be a B-cell epitope or not. Although tools belonging to the first category can also be executed for small peptides, it would be unjustifiable to compare them with the tools belonging to the second category. For example, Bepipred-3.0, Bepipred-2.0, Bepipred-1.0, have excellent predictive capabilities when tested on whole protein sequences but they do not perform in a desired manner when tested on short peptides, as observed in a recent study (da Silva, Ascher et al. 2023). Our BLMPred model falls into the second category and here, we conduct a detailed comparative performance analysis of BLMPred with the freely available standalone tools belonging to the same category that we were able to install and execute – SVMTriP, LBEEP, and epitope1D. For SVMTriP, six separate models were trained separately on epitopes of length 10, 12, 14, 16, 18, and 20 amino acids (Yao, Zheng et al. 2020). We selected the reportedly best-performing SVMTriP model trained on epitopes of 20 amino acids in length. LBEEP, SVMTriP, epitope1D, only accept as input peptides within the length ranges 6-15, 10-20, and 6-, respectively. Although BLMPred models do not have any strict length-based restrictions and can be executed for any length peptide, they have been trained and thus perform best on peptides whose length is in the range of 5-60. Hence, we utilized different groups of samples from the BLMPred_benchmark dataset to fulfill the length-based restrictions of the selected tools for an unbiased detailed comparative analysis of their performance (Table 1).

**Table 1.** Comparison of BLMPred (name marked in bold font) with the previously published methods. The best achieved values for each of the performance metrics are highlighted in bold font. Lowest achieved values for FP and FN are in bold italics font.

| Model |  | Sample selection from BLMPred_benchmark dataset |  |  | Performance metrics |  |  |  |  |  |  |  |  |  |  |
| --- | --- | --- | --- | --- | --- | --- | --- | --- | --- | --- | --- | --- | --- | --- | --- |
| Name | Length-based restriction (min-max) | Length range | #Epitopes | #Non-epitopes | Accuracy | Precision | Recall | F1 score | Specificity | AUROC | AUPRC | TP | FP | TN | FN |
| **BLMPred** | NA | 6-71 | 2928 | 1000 | **0.72** | 0.86 | **0.75** | **0.80** | 0.64 | **0.69** | **0.83** | **2194 (74.9%)** | 361 (36.1%) | 639 (63.9%) | **734 (25.1%)** |
| epitope1D | 6- |  |  |  | 0.31 | **0.89** | 0.09 | 0.16 | **0.97** | 0.53 | 0.76 | 259 (8.8%) | **31 (3.1%)** | **969 (96.9%)** | 2669 (91.2%) |
| **BLMPred** | NA | 20 | 188 | 41 | **0.66** | 0.81 | **0.76** | **0.78** | 0.17 | 0.46 | 0.81 | **143 (76%)** | 34 (83%) | 7 (17%) | **45 (24%)** |
| epitope1D | 6- |  |  |  | 0.21 | **1.00** | 0.03 | 0.06 | **1.00** | **0.52** | **0.83** | 6 (3.2%) | **0 (0%)** | **41 (100%)** | 182 (96.8%) |
| SVMTriP | 20 |  |  |  | 0.30 | 0.78 | 0.21 | 0.33 | 0.72 | 0.46 | 0.82 | 39 (20.7%) | 11 (31.7%) | 28 (68.3%) | 149 (79.3%) |
| **BLMPred** | NA | 6-15 | 546 | 400 | **0.63** | 0.78 | **0.50** | **0.60** | 0.80 | **0.65** | **0.68** | **273 (50%)** | 79 (19.8%) | 321 (80.2%) | **273 (50%)** |
| LBEEP | 6-15 |  |  |  | 0.47 | 0.56 | 0.35 | 0.43 | 0.63 | 0.49 | 0.57 | 192 (35.2%) | 149 (37%) | 252 (63%) | 355 (64.8%) |
| epitope1D | 6- |  |  |  | 0.45 | **0.90** | 0.05 | 0.09 | **0.99** | 0.52 | 0.59 | 26 (4.8%) | **3 (0.5%)** | **398 (99.5%)** | 521 (95.2%) |

As shown in Table 1, BLMPred correctly identifies 75% 50%, and 76% of the epitopes in the length ranges of 6-71, 6-15, and 20, respectively, in the benchmarking dataset. Also, BLMPred correctly predicts 64% and 80%, non-B-cell-epitopes in the length ranges of 6-71 and 6-15, respectively. Although the model identifies 143 epitopes out of 188 epitopes, it fails to identify most of the non-B-cell-epitopes, resulting in a low specificity for the 20-length range. We think that this problem happens because the model was trained on very few samples of length 20. Only 1.6% of our training dataset consisted of peptides with the length of exactly 20aa. Based on this performance analysis, we can suggest that BLMPred has a good capability of predicting epitopes from a collection of peptides and it outperforms other tools selected for the comparative study, especially for the peptide length ranges of 6-71 and 6-15.

## 4 Conclusions

The existing machine learning-based B-cell epitope prediction tools can be broadly partitioned into two categories. One category including tools such as Bepipred-3.0, Bepipred-2.0, epiDope, are trained on complete antigen protein sequences, whereas the other category including tools such as SVMTriP, LBEEP, epitope1D, are trained on exact linear B-cell epitope sequences. The first category seems to perform poorly on short peptides as observed in a recent study (da Silva, Ascher et al. 2023). Both categories are equally important for antibody epitope predictions. The first category is mainly beneficial when the goal is to identify all B-cell epitopes from an entire protein sequence. The second category is applicable when the user seeks to determine if a specific peptide could serve as a potential B-cell epitope. Previous reports suggest that epitope prediction from conserved protein sequences derived from Multiple Sequence Alignments (MSAs) is more accurate (Yasmin and Nabi 2016). Peptide-based B-cell epitope prediction tools are particularly well suitable for classifying conserved MSA blocks as antibody epitopes or not.

BLMPred is a simple binary classifier developed by training a Support Vector Machine on a huge non-redundant experimentally validated dataset composed of 222030 peptides numerically embedded by the ProtTrans pLM. The dataset has been carefully constructed by maintaining uniform length distributions among both positive and negative samples collected from the IEDB database. A 10-fold cross-validation revealed that the BLMPred model outperformed other traditional machine learning algorithms. The BLMPred model is characterized by low rates of both type-I and type-II errors and therefore has a good predictive capability of identifying B-cell epitopes and the non-antibody epitopes, as evident from Figure 2. BLMPred outperforms other tested tools on all partitions of the benchmarking datasets in terms of accuracy, recall, F1 score, MCC, AUROC, and AUPRC values (Table 1).

The BLMPred model is available as a Github repository (https://github.com/bdbarnalidas/BLMPred.git) with thorough instructions for the users and can be easily cloned/downloaded and executed by the experts and non-experts alike. The B-cell epitopes predicted by BLMPred can be utilized in the fields of immunology and biotechnology for vaccine development and antibody engineering. As a future vision, we can anticipate that combining structure-based embeddings and sequence-based embeddings may further improve the predictive potential of BLMPred. We also wish to extend our approach by training BLMPred on entire protein sequences embedded by ProtTrans to predict one or multiple B-cell epitopes from the protein sequences. We are planning to apply both per-residue as well as per-protein-based embeddings for that purpose. We can also test other widely used pLMs like ESM (Lin, Akin et al. 2023). Another probable future peptide-based approach would be to map peptides to protein sequences, generate embeddings for proteins, extract sub-embeddings corresponding to the peptide of interest, and use them for training machine learning models. We anticipate that this approach may improve the overall performance since pLMs the embedding vectors for entire protein sequence capture long-range dependencies possibly making the resulting representation more informative.

## Declarations

## Abbreviations

Not applicable

## Ethics approval and consent to participate

Not applicable

## Consent for publication

All authors consent for publication.

## Availability of data and materials

We used the experimentally validated linear B-cell epitopes from the IEDB database (https://www.iedb.org). The BLMPred datasets created for this paper are publicly available at https://github.com/bdbarnalidas/BLMPred/tree/main/BLMPred_Datasets. The benchmarking datasets used for a performance assessment between different models are publicly available at https://github.com/bdbarnalidas/BLMPred/tree/main/BLMPred_Datasets.

## Authors’ contributions

B.D.: Conception, Design, Data acquisition, Analysis, Software, Writing original draft; D.F.: Conception, Analysis, Supervision, Funding acquisition, Review manuscript.

## Acknowledgements

Not applicable

