## Supplementary file for "BLMPred: predicting linear B-cell epitopes using pre-trained protein language models and machine learning"

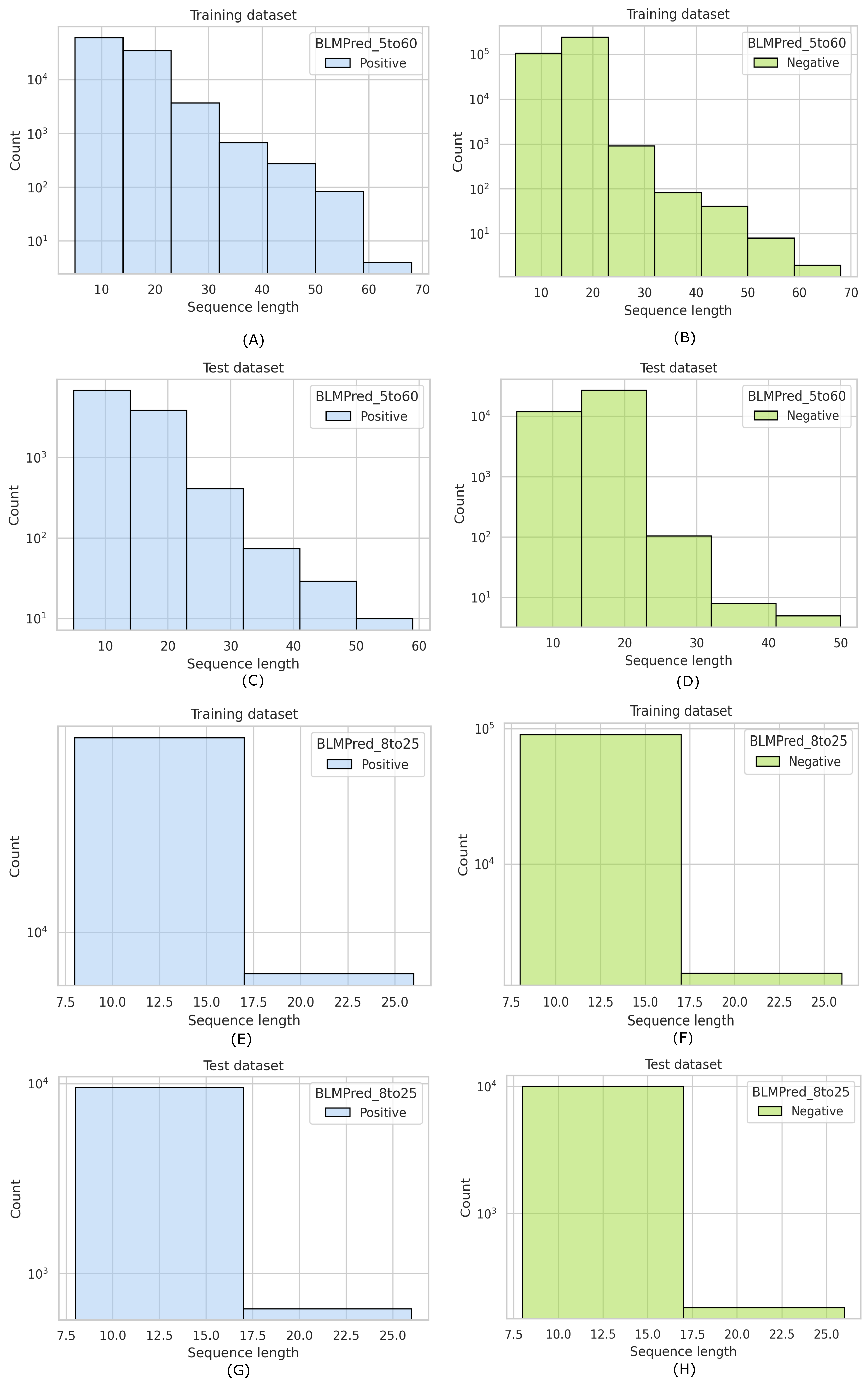

**Figure S1.** Sequence length distributions. (A) Positive samples in BLMPred_5to60_training dataset, (B) Negative samples in BLMPred_5to60_training dataset, (C) Positive samples in BLMPred_5to60_test dataset, (D) Negative samples in BLMPred_5to60_test dataset, (E) Positive samples in BLMPred_8to25_training dataset, (F) Negative samples in BLMPred_8to25_training dataset, (G) Positive samples in BLMPred_8to25_test dataset, (H) Negative samples in BLMPred_8to25_test dataset.

**Supplementary Table S1.** Cross-validation performance of models trained on BLMPred_5to60_training dataset. For each classifier, mean (standard deviation) of the performance metrics for 10-fold cross-validation are shown. Maximum values of the performance metrics and the best trained model are highlighted in bold.

| Dataset/Model | BLMPred_5to60 | | | | | | | |
| --- | --- | --- | --- | --- | --- | --- | --- | --- |
|  | Accuracy | Precision | Recall | F1 score | Specificity | MCC | AUROC | AUPRC |
| AdaBoost classifier | 0.686 (0.0032) | 0.691 (0.0047) | 0.670 (0.0049) | 0.681 (0.0039) | 0.701 (0.0039) | 0.371 (0.0063) | 0.686 (0.0032) | 0.628 (0.0044) |
| Bagging classifier | 0.763 (0.0031) | 0.800 (0.0049) | 0.701 (0.0049) | 0.747 (0.0039) | 0.825 (0.0045) | 0.529 (0.0064) | 0.763 (0.0032) | 0.710 (0.0049) |
| Extra trees classifier | 0.769 (0.0016) | 0.786 (0.0026) | 0.739 (0.0025) | 0.762 (0.0016) | 0.798 (0.0031) | 0.538 (0.0032) | 0.769 (0.0016) | 0.711 (0.0022) |
| Gaussian Naïve Bayes | 0.662 (0.0038) | 0.661 (0.0039) | 0.665 (0.0059) | 0.663 (0.0044) | 0.659 (0.0050) | 0.325 (0.0077) | 0.662 (0.0038) | 0.607 (0.0036) |
| Histogram-based gradient boosting classifier | 0.745 (0.0023) | 0.767 (0.0049) | 0.704 (0.0036) | 0.734 (0.0027) | 0.786 (0.0045) | 0.492 (0.0047) | 0.745 (0.0023) | 0.688 (0.0041) |
| k-nearest neighbors | 0.789 (0.0020) | 0.799 (0.0022) | 0.773 (0.0041) | 0.786 (0.0022) | 0.807 (0.0025) | 0.579 (0.0039) | 0.789 (0.0020) | 0.732 (0.0021) |
| Linear discriminant analysis | 0.725 (0.0024) | 0.735 (0.0044) | 0.704 (0.0046) | 0.719 (0.0028) | 0.746 (0.0049) | 0.450 (0.0048) | 0.725 (0.0024) | 0.665 (0.0036) |
| Logistic regression | 0.726 (0.0017) | 0.734 (0.0040) | 0.708 (0.0044) | 0.721 (0.0030) | 0.743 (0.0036) | 0.451 (0.0034) | 0.726 (0.0017) | 0.666 (0.0039) |
| Multi-layer perceptron | 0.777 (0.0036) | 0.776 (0.0176) | 0.781 (0.0280) | 0.778 (0.0065) | 0.773 (0.0315) | 0.555 (0.0071) | 0.777 (0.0038) | 0.715 (0.0074) |
| Quadratic discriminant analysis | 0.742 (0.0034) | 0.764 (0.0047) | 0.701 (0.0056) | 0.731 (0.0031) | 0.784 (0.0057) | 0.486 (0.0067) | 0.742 (0.0033) | 0.685 (0.0031) |
| Random forest | 0.774 (0.0018) | 0.797 (0.0039) | 0.735 (0.0041) | 0.765 (0.0027) | 0.813 (0.0028) | 0.550 (0.0036) | 0.774 (0.0019) | 0.719 (0.0038) |
| **Support vector machine** | **0.839** (0.0027) | **0.849** (0.0046) | **0.824** (0.0037) | **0.837** (0.0033) | **0.855** (0.0041) | **0.679** (0.0055) | **0.839** (0.0028) | **0.788** (0.0047) |
| XGBoost | 0.754 (0.0029) | 0.771 (0.0038) | 0.723 (0.0048) | 0.746 (0.0034) | 0.785 (0.0028) | 0.509 (0.0058) | 0.754 (0.0029) | 0.696 (0.0038) |
| Explainable Boosting classifier | 0.741 (0.0029) | 0.755 (0.0041) | 0.714 (0.0061) | 0.734 (0.0031) | 0.768 (0.0046) | 0.483 (0.0055) | 0.741 (0.0027) | 0.682 (0.0034) |

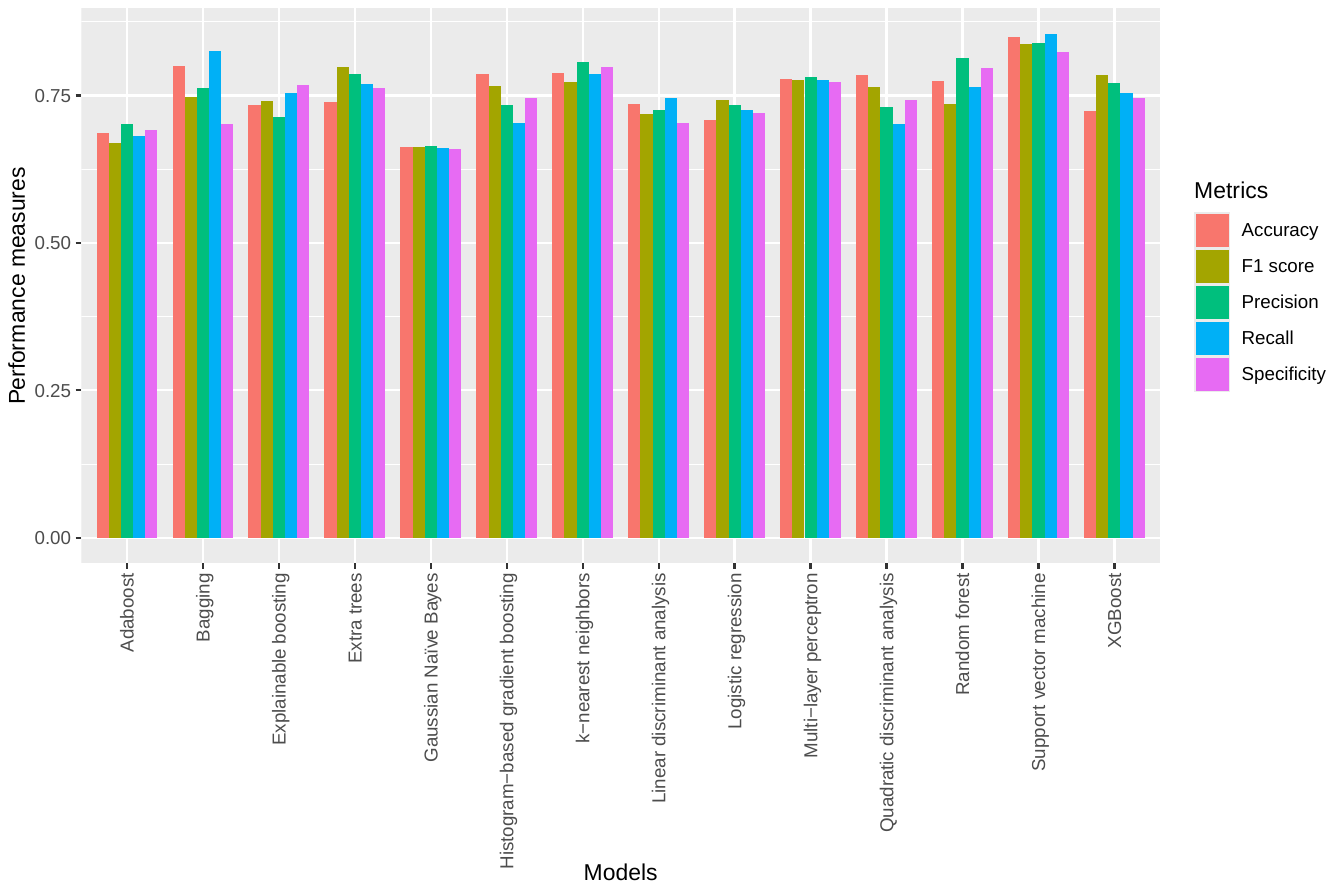

**Figure S2.** Cross-validation performance of the models trained on the BLMPred_5to60_training dataset. For each classifier, mean values of the performance metrics for 10-fold cross-validated models are shown.

**Supplementary Table S2.** Performance of models trained on BLMPred_5to60_training dataset on the blind BLMPred_5to60_test dataset. For each classifier, mean (standard deviation) of the performance metrics for 10 models trained in each fold of 10-fold cross-validation when applied on the blind test dataset are shown. Maximum values of the performance metrics and the best performing model are highlighted in bold.

| Dataset/Model | BLMPred_5to60 | | | | | | | |
| --- | --- | --- | --- | --- | --- | --- | --- | --- |
|  | Accuracy | Precision | Recall | F1 score | Specificity | MCC | AUROC | AUPRC |
| AdaBoost classifier | 0.686 (0.0014) | 0.690 (0.0018) | 0.673 (0.0022) | 0.682 (0.0014) | 0.698 (0.0028) | 0.371 (0.0028) | 0.686 (0.0014) | 0.628 (0.0013) |
| Bagging classifier | 0.765 (0.0007) | 0.803 (0.0011) | 0.704 (0.0024) | 0.749 (0.0011) | 0.827 (0.0016) | 0.535 (0.0014) | 0.765 (0.0007) | 0.713 (0.0007) |
| Extra trees classifier | 0.770 (0.0014) | 0.786 (0.0022) | 0.741 (0.0026) | 0.763 (0.0015) | 0.799 (0.0029) | 0.541 (0.0028) | 0.770 (0.0014) | 0.712 (0.0016) |
| Gaussian Naïve Bayes | 0.663 (0.0007) | 0.661 (0.0010) | 0.666 (0.0007) | 0.664 (0.0005) | 0.659 (0.0019) | 0.325 (0.0015) | 0.663 (0.0007) | 0.607 (0.0007) |
| Histogram-based gradient boosting classifier | 0.746 (0.0007) | 0.766 (0.0010) | 0.707 (0.0017) | 0.735 (0.0089) | 0.785 (0.0015) | 0.493 (0.0014) | 0.746 (0.0073) | 0.688 (0.0007) |
| k-nearest neighbors | 0.790 (0.0008) | 0.799 (0.0011) | 0.775 (0.0014) | 0.787 (0.0008) | 0.806 (0.0014) | 0.581 (0.0016) | 0.790 (0.0008) | 0.732 (0.0009) |
| Linear discriminant analysis | 0.727 (0.0004) | 0.737 (0.0005) | 0.706 (0.0009) | 0.721 (0.0005) | 0.748 (0.0007) | 0.454 (0.0009) | 0.727 (0.0004) | 0.667 (0.0004) |
| Logistic regression | 0.726 (0.0008) | 0.734 (0.0009) | 0.709 (0.0013) | 0.721 (0.0008) | 0.744 (0.0011) | 0.453 (0.0015) | 0.726 (0.0007) | 0.666 (0.0007) |
| Multi-layer perceptron | 0.777 (0.0047) | 0.778 (0.0188) | 0.779 (0.0294) | 0.778 (0.0068) | 0.776 (0.0330) | 0.556 (0.0087) | 0.777 (0.0047) | 0.716 (0.0078) |
| Quadratic discriminant analysis | 0.746 (0.0007) | 0.769 (0.0023) | 0.704 (0.0027) | 0.735 (0.0007) | 0.789 (0.0035) | 0.494 (0.0015) | 0.746 (0.0007) | 0.689 (0.0010) |
| Random forest | 0.777 (0.0007) | 0.799 (0.0010) | 0.740 (0.0011) | 0.768 (0.0008) | 0.814 (0.0012) | 0.555 (0.0015) | 0.777 (0.0007) | 0.721 (0.0008) |
| **Support vector machine** | **0.841** (0.0010) | **0.853** (0.0012) | **0.823** (0.0013) | **0.838** (0.0011) | **0.859** (0.0013) | **0.682** (0.0021) | **0.841** (0.0010) | **0.791** (0.0013) |
| XGBoost | 0.756 (0.0017) | 0.773 (0.0017) | 0.725 (0.0030) | 0.748 (0.0019) | 0.787 (0.0020) | 0.513 (0.0033) | 0.756 (0.0017) | 0.698 (0.0016) |
| Explainable Boosting classifier | 0.743 (0.0011) | 0.756 (0.0013) | 0.717 (0.0015) | 0.736 (0.0012) | 0.768 (0.0014) | 0.486 (0.0023) | 0.743 (0.0011) | 0.638 (0.0011) |

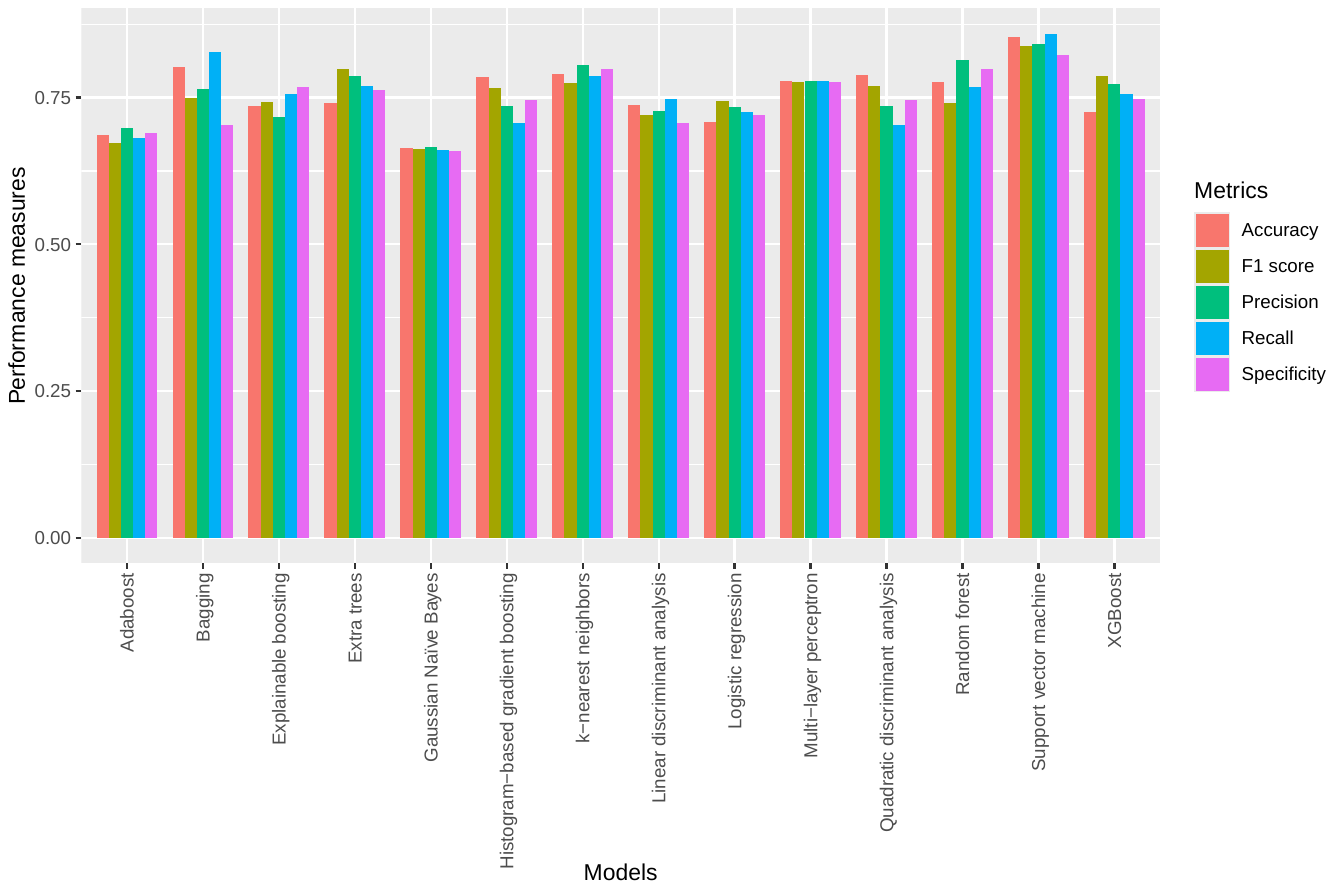

**Figure S3.** Performance of models trained on BLMPred_5to60_training dataset on the blind BLMPred_5to60_test dataset. For each classifier, mean values of the performance metrics for 10 models trained in each fold of 10-fold cross-validation when applied on the blind test dataset are shown.

**Supplementary Table S3.** Cross-validation performance of models trained on BLMPred_8to25_training dataset. For each classifier, mean (standard deviation) of the performance metrics for 10-fold cross-validation are shown. Maximum values of the performance metrics and the best trained model are highlighted in bold.

| Dataset/Model | BLMPred_8to25 | | | | | | | |
| --- | --- | --- | --- | --- | --- | --- | --- | --- |
|  | Accuracy | Precision | Recall | F1 score | Specificity | MCC | AUROC | AUPRC |
| AdaBoost classifier | 0.675 (0.0044) | 0.679 (0.0054) | 0.665 (0.0078) | 0.672 (0.0058) | 0.686 (0.0037) | 0.351 (0.0089) | 0.675 (0.0045) | 0.619 (0.0054) |
| Bagging classifier | 0.757 (0.0043) | 0.790 (0.0057) | 0.699 (0.0061) | 0.742 (0.0052) | 0.814 (0.0039) | 0.517 (0.0086) | 0.757 (0.0043) | 0.703 (0.0058) |
| Extra trees classifier | 0.758 (0.0024) | 0.771 (0.0031) | 0.733 (0.0036) | 0.752 (0.0023) | 0.782 (0.0036) | 0.516 (0.0047) | 0.758 (0.0023) | 0.699 (0.0026) |
| Gaussian Naïve Bayes | 0.633 (0.0036) | 0.617 (0.0051) | 0.702 (0.0029) | 0.657 (0.0035) | 0.565 (0.0063) | 0.269 (0.0068) | 0.633 (0.0034) | 0.582 (0.0043) |
| Histogram-based gradient boosting classifier | 0.736 (0.0029) | 0.753 (0.0055) | 0.703 (0.0046) | 0.727 (0.0035) | 0.769 (0.0059) | 0.473 (0.0059) | 0.736 (0.0029) | 0.678 (0.0048) |
| k-nearest neighbors | 0.782 (0.0029) | 0.789 (0.0051) | 0.769 (0.0031) | 0.779 (0.0028) | 0.794 (0.0054) | 0.564 (0.0059) | 0.782 (0.0029) | 0.723 (0.0044) |
| Linear discriminant analysis | 0.717 (0.0039) | 0.723 (0.0049) | 0.702 (0.0061) | 0.713 (0.0041) | 0.731 (0.0046) | 0.434 (0.0079) | 0.717 (0.0039) | 0.657 (0.0043) |
| Logistic regression | 0.716 (0.0028) | 0.721 (0.0049) | 0.704 (0.0050) | 0.713 (0.0038) | 0.728 (0.0039) | 0.432 (0.0056) | 0.716 (0.0028) | 0.656 (0.0046) |
| Multi-layer perceptron | 0.772 (0.0025) | 0.778 (0.0197) | 0.766 (0.0333) | 0.771 (0.0078) | 0.779 (0.0348) | 0.546 (0.0053) | 0.772 (0.0026) | 0.712 (0.0071) |
| Quadratic discriminant analysis | 0.699 (0.0031) | 0.671 (0.0048) | 0.782 (0.0046) | 0.722 (0.0037) | 0.618 (0.0057) | 0.405 (0.0059) | 0.699 (0.0029) | 0.634 (0.0046) |
| Random forest | 0.765 (0.0035) | 0.784 (0.0035) | 0.732 (0.0043) | 0.757 (0.0037) | 0.798 (0.0035) | 0.532 (0.0069) | 0.765 (0.0035) | 0.708 (0.0037) |
| **Support vector machine** | **0.833** (0.0023) | **0.848** (0.0026) | **0.811** (0.0042) | **0.829** (0.0027) | **0.854** (0.0016) | **0.666** (0.0044) | **0.833** (0.0022) | **0.782** (0.0032) |
| XGBoost | 0.747 (0.0026) | 0.760 (0.0039) | 0.721 (0.0042) | 0.740 (0.0033) | 0.773 (0.0032) | 0.495 (0.0052) | 0.747 (0.0026) | 0.688 (0.0042) |
| Explainable Boosting classifier | 0.733 (0.0039) | 0.744 (0.0053) | 0.710 (0.0040) | 0.727 (0.0038) | 0.756 (0.0066) | 0.467 (0.0079) | 0.733 (0.0039) | 0.674 (0.0045) |

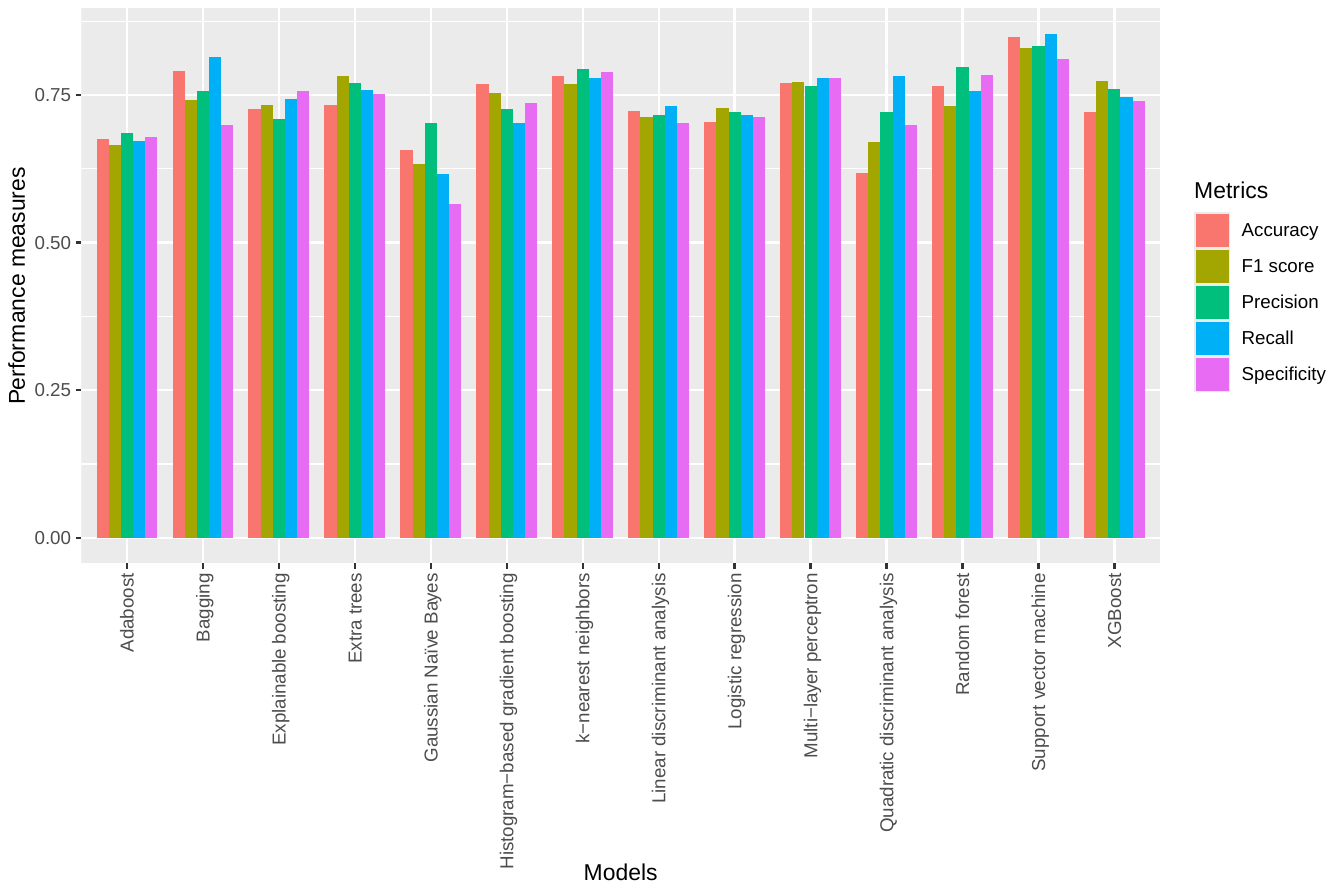

**Figure S4.** Cross-validation performance of the models trained on the BLMPred_8to25_training dataset. For each classifier, mean values of the performance metrics for 10-fold cross-validated models are shown.

**Supplementary Table S4.** Performance of models trained on BLMPred_8to25_training dataset on the blind BLMPred_8to25_test dataset. For each classifier, mean (standard deviation) of the performance metrics for 10 models trained in each fold of 10-fold cross-validation when applied on the blind test dataset are shown. Maximum values of the performance metrics and the best performing model are highlighted in bold.

| Dataset/Model | BLMPred_8to25 | | | | | | | |
| --- | --- | --- | --- | --- | --- | --- | --- | --- |
|  | Accuracy | Precision | Recall | F1 score | Specificity | MCC | AUROC | AUPRC |
| AdaBoost classifier | 0.672 (0.0014) | 0.675 (0.0018) | 0.665 (0.0023) | 0.669 (0.0014) | 0.679 (0.0029) | 0.344 (0.0027) | 0.672 (0.0014) | 0.616 (0.0012) |
| Bagging classifier | 0.749 (0.0013) | 0.782 (0.0013) | 0.692 (0.0021) | 0.734 (0.0015) | 0.807 (0.0012) | 0.502 (0.0025) | 0.749 (0.0013) | 0.695 (0.0013) |
| Extra trees classifier | 0.752 (0.0018) | 0.765 (0.0022) | 0.729 (0.0023) | 0.747 (0.0018) | 0.776 (0.0026) | 0.506 (0.0035) | 0.752 (0.0018) | 0.693 (0.0019) |
| Gaussian Naïve Bayes | 0.632 (0.0008) | 0.615 (0.0009) | 0.704 (0.0006) | 0.657 (0.0004) | 0.559 (0.0021) | 0.267 (0.0016) | 0.632 (0.0008) | 0.581 (0.0006) |
| Histogram-based gradient boosting classifier | 0.731 (0.0011) | 0.747 (0.0017) | 0.699 (0.0017) | 0.723 (0.0011) | 0.763 (0.0022) | 0.464 (0.0023) | 0.731 (0.0011) | 0.673 (0.0012) |
| k-nearest neighbors | 0.777 (0.0009) | 0.786 (0.0009) | 0.761 (0.0021) | 0.773 (0.0011) | 0.793 (0.0014) | 0.554 (0.0017) | 0.777 (0.0009) | 0.718 (0.0009) |
| Linear discriminant analysis | 0.712 (0.0008) | 0.717 (0.0009) | 0.700 (0.0010) | 0.708 (0.0008) | 0.723 (0.0014) | 0.423 (0.0016) | 0.712 (0.0008) | 0.652 (0.0008) |
| Logistic regression | 0.713 (0.0008) | 0.717 (0.0009) | 0.705 (0.0006) | 0.711 (0.0007) | 0.721 (0.0012) | 0.426 (0.0016) | 0.713 (0.0008) | 0.653 (0.0008) |
| Multi-layer perceptron | 0.769 (0.0025) | 0.775 (0.0190) | 0.763 (0.0332) | 0.768 (0.0078) | 0.777 (0.0339) | 0.541 (0.0050) | 0.7769 (0.0025) | 0.709 (0.0059) |
| Quadratic discriminant analysis | 0.696 (0.0011) | 0.668 (0.0015) | 0.779 (0.0017) | 0.719 (0.0006) | 0.613 (0.0033) | 0.398 (0.0019) | 0.696 (0.0011) | 0.631 (0.0010) |
| Random forest | 0.761 (0.0008) | 0.778 (0.0010) | 0.731 (0.0012) | 0.753 (0.0009) | 0.791 (0.0012) | 0.523 (0.0016) | 0.761 (0.0008) | 0.703 (0.0009) |
| **Support vector machine** | **0.829** (0.0009) | **0.845** (0.0018) | **0.807** (0.0009) | **0.825** (0.0008) | **0.852** (0.0021) | **0.659** (0.0019) | **0.829** (0.0009) | **0.778** (0.0014) |
| XGBoost | 0.743 (0.0017) | 0.756 (0.0018) | 0.717 (0.0034) | 0.736 (0.0021) | 0.769 (0.0023) | 0.487 (0.0034) | 0.743 (0.0017) | 0.684 (0.0016) |
| Explainable Boosting classifier | 0.726 (0.0009) | 0.735 (0.0015) | 0.706 (0.0011) | 0.720 (0.0009) | 0.745 (0.0020) | 0.452 (0.0020) | 0.726 (0.0009) | 0.666 (0.0011) |

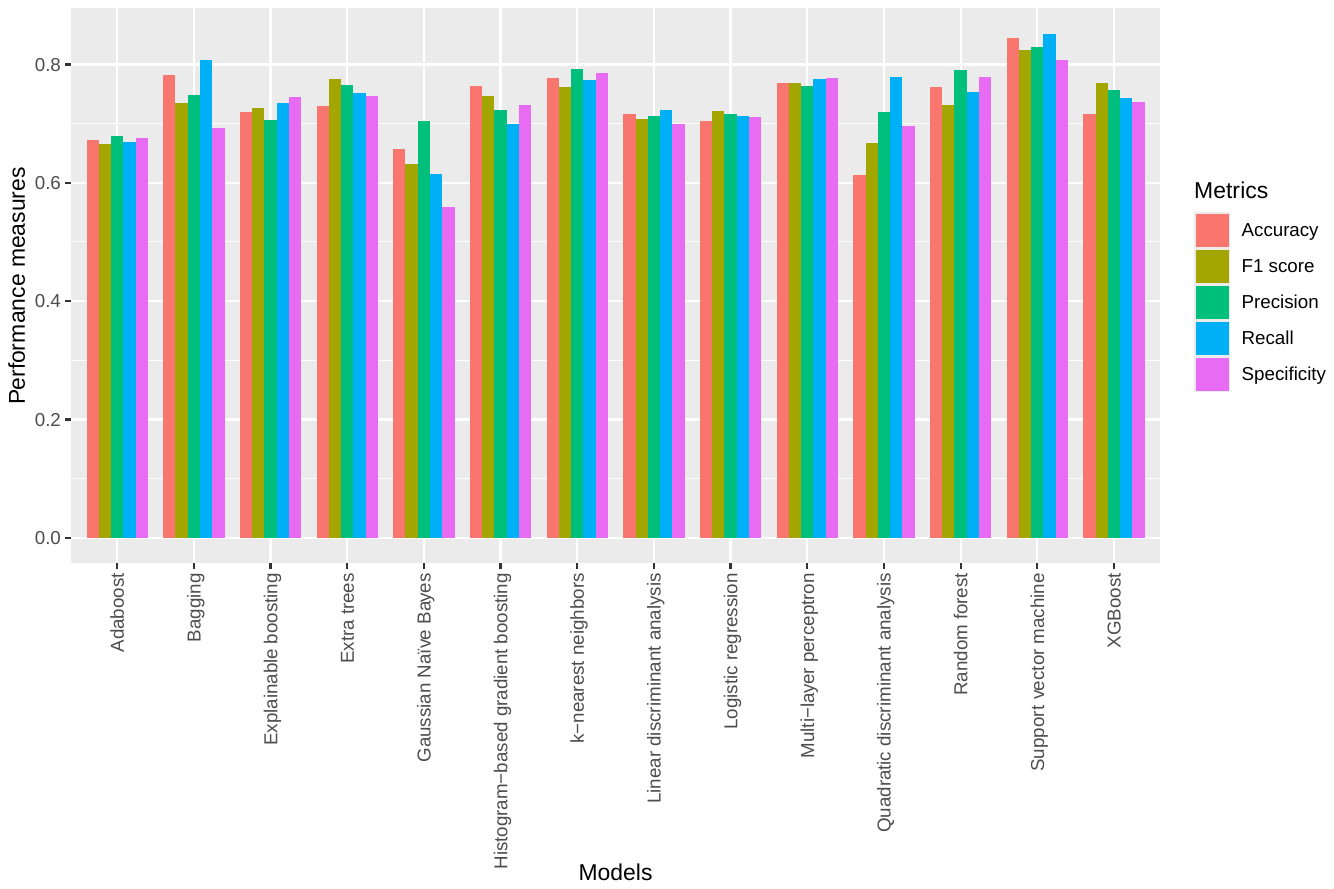

**Figure S5.** Performance of the models trained on the BLMPred_8to25_training dataset on the blind BLMPred_8to25_test dataset. For each classifier, mean values of the performance metrics for 10 models trained in each fold of 10-fold cross-validation when applied on the blind test dataset are shown.

**Supplementary Table S5.** Detailed summary of the computational tools considered in our comparative study. Performance metrics of the tools as reported in their respective publications are presented, with the best values highlighted in bold. Our models are also highlighted in bold.

| Model | Algorithm | Dataset | Features | Reported performance | | | | | | | |
| --- | --- | --- | --- | --- | --- | --- | --- | --- | --- | --- | --- |
|  |  |  |  | Accuracy | Precision | Recall | MCC | Specificity | F1 score | AUROC | AUPRC |
| **BLMPred_5to60** | Support Vector Machine | Linear BCE from IEDB | ProtTrans embeddings | **0.841** | **0.853** | **0.823** | 0.682 | **0.859** | **0.838** | 0.841 | **0.791** |
| **BLMPred_8to25** |  |  |  | 0.829 | 0.845 | 0.807 | 0.659 | 0.852 | 0.825 | 0.829 | 0.778 |
| SVMTriP | Support Vector Machine | Linear BCE from IEDB | Tri-peptide similarity and propensity scores | NA | 0.552 | 0.801 | NA | NA | 0.693 | 0.702 | NA |
| LBEEP | Support Vector Machine & AdaBoost Random Forest | Linear BCE | Amino acid composition-based feature | 0.73 | NA | NA | NA | NA | NA | NA | NA |
| epitope1D | Random Forest and Explainable Boosting Machine | Linear BCE from IEDB | Graph-based signature representation of protein sequences and  organism ontology information | NA | NA | NA | **0.72** | NA | NA | **0.93** | NA |
